# Fungal susceptibility and early flowering in pennycress (*Thlaspi arvense*) are conferred by naturally occurring mutations in histone demethylase Jumonji 14

**DOI:** 10.1101/2025.05.14.654108

**Authors:** Jennette M. Codjoe, Alice Kujur, Jammi Prasanthi Sirasani, Anastasia Shamin, Thomas Sauer, Krishan Rai, Tim Ulmasov, Ratan Chopra, Dilip M Shah

**Affiliations:** Donald Danforth Plant Science Center, St. Louis, MO 63132, USA; CoverCress Inc., St. Louis, MO 63132, USA

**Keywords:** pennycress, early flowering, disease susceptibility, *Sclerotinia sclerotiorum*, *Alternaria japonica*, Jumonji 14, histone demethylase, winter oilseed, pattern-triggered immunity

## Abstract

Pennycress (*Thlaspi arvense*) is a winter oilseed domesticated recently to be incorporated as an intermediate crop between the existing cropping systems of the US Midwest. We show that a natural accession of pennycress, 2032, is more susceptible to the necrotrophic fungal pathogens *Sclerotinia sclerotiorum* and *Alternaria japonica* than the reference pennycress accession MN106. A previously identified marker associated with early flowering and maturity in pennycress was found to be present in a gene homologous to Arabidopsis *Jumonji 14 (JMJ14).* It has been reported that AtJMJ14 promotes disease resistance and represses flowering, and greenhouse studies of breeding populations confirmed this phenomenon in pennycress. Plants with the 2032 *TaJMJ14* allele were more susceptible to fungi and flowered early. CRISPR-Cas9 editing was used to generate additional *TaJMJ14* alleles. A 9-base pair deletion in the 6^th^ exon of *TaJMJ14* showed trends of early flowering and *S. sclerotiorum* susceptibility, whereas a complete loss-of-function allele led to infertility. We further investigated the transcriptomes of MN106 and 2032 plants in the early stages of *S. sclerotiorum* and *A. japonica* infection to identify potential resistance and susceptibility genes. Differences in the expression of pathogen-associated molecular pattern-triggered immunity (PTI)-associated genes led us to discover that 2032 plants have defects in elicitor-triggered oxidative bursts. The transcriptional responses unique to each accession lay a foundation for future gene-editing and breeding approaches to keep the beneficial early flowering phenotype conferred by 2032 but uncouple it from disease susceptibility.

## INTRODUCTION

Pennycress (*Thlaspi arvense*) is being domesticated as a cash cover crop whose oil-rich seeds can be used for biofuel production (Phippen et al., 2022). In the Midwest of the United States, pennycress can be grown during the winter in between corn and soybean rotations. This has the potential to increase productivity of these farms by growing another crop on what is otherwise fallow land (Sedbrook et al., 2014). In CoverCress Inc. research fields in the Midwest, pennycress predominantly becomes infected with two fungal diseases: white mold and Alternaria black spot (Kujur and Codjoe et al., 2025). These diseases have the potential to impact pennycress yield as they do in other crops (Peltier et al., 2012; Al-lami et al., 2019; O’Sullivan et al., 2021), although this has not been studied.

White mold disease is caused by the fungus *Sclerotinia sclerotiorum,* which infects a wide variety of dicot plants (Bolton et al., 2006). Alternaria black spot (also called leaf spot) disease occurs specifically in Brassica species and is caused by *Alternaria brassicae, A. brassicicola, A. raphani,* or *A. japonica* (Nowicki et al., 2012; Al-lami et al., 2020). *A. brassicicola* and *A. raphani* have been reported to infect wild growing pennycress (Cobb and Dillard, 1998), and we recently found that *A. japonica* can also infect pennycress research fields (Kujur and Codjoe et al., 2025). Both *S. sclerotiorum* and *Alternaria spp.* are necrotrophic pathogens, meaning that they kill host cells to feed on dead or dying cells (Doehlemann et al., 2017). In other plants, resistance to white mold and black spot is governed by quantitative resistance genes, many of which function in pathogen-associated molecular pattern-triggered immunity (PTI). PTI is an innate immune response triggered by cell-surface pathogen perception, beginning a signaling cascade that leads to transcriptional changes and the production of hormones and secondary metabolites that limit pathogen growth (Wang et al., 2019; Singh et al., 2021; Duo et al., 2025). Putative resistance genes for powdery mildew have recently been identified in pennycress (Galanti et al., 2024); however, the genetic sources of resistance (and susceptibility) to *S. sclerotiorum* and *A. japonica* are unknown.

Epigenetic regulation, including the post-translational modification of histones, is one way that transcription is reprogrammed as plants adaptively respond to pathogens. Certain histone modifications, like H3K4 and H3K36 methylation remodel the chromatin to favor gene expression, whereas others, like H3K9 and H3K27 methylation repress transcription (Zhi and Chang, 2021). Histone demethylation by Jumonji C (JmjC) domain-containing proteins has previously been implicated in the response to various pathogens. Arabidopsis Jumonji 14 (JMJ14) and JMJ27 positively regulate defense against the bacterial pathogen *Pseudomonas syringae* (Li et al., 2020; Dutta et al., 2017), and rice JMJ704 and JMJ705 promote resistance to *Xanthomonas oryzae* (Li et al., 2013; Hou et al., 2015). In all these cases, resistance is presumably provided by the induction of resistance genes or repression of negative defense regulators, mediated directly or indirectly by JMJ demethylation activity. Disease-related genes are not the only targets; JMJ14 and JMJ27 both regulate the expression of genes controlling flowering, and both are negative regulators of flowering time (Jeong et al., 2009; Lu et al., 2010; Yang et al., 2010; Dutta et al., 2017). JMJ14 regulates flowering by removing transcription-activating H3K4 trimethylation at *Flowering Locus T (FT)* chromatin, repressing *FT* expression and hence flowering (Jeong et al., 2009). Early flowering genes are attractive targets for pennycress domestication, as early maturity allows for the timely harvest of pennycress in the spring without significantly delaying the planting of the subsequent crop (Chopra et al., 2020; Basnet and Ellison, 2024).

In this study we characterized the earliness and disease susceptibility of two natural accessions of pennycress from North America, MN106 and 2032. MN106 was isolated near Coates, MN, USA and is the source of the *T. arvense* v2 reference genome (Dorn et al. 2013; Nunn et al. 2020). 2032 is a proprietary CoverCress line from CO, USA that has considerable phenotypic and genetic differences compared to MN106. Here we report that 2032 plants flower earlier and are more susceptible to white mold and Alternaria black spot than MN106 plants due to a presumed loss-of-function mutation in the *TaJMJ14* gene. Using comparative transcriptomics of these accessions during infection, we identified putative resistance and susceptibility genes that may underly their differences in susceptibility to fungal diseases.

## RESULTS

### Mutations present in the TaJMJ14 gene of 2032 plants co-segregate with early flowering and maturity phenotypes

A molecular marker was identified at CoverCress Inc. that associated with earliness, a critical trait in its breeding program. This marker was a single base pair insertion in an unannotated region of chromosome 7 of the *T. arvense* v2 genome. A BLASTn search of this region against the Arabidopsis genome found that it contained a sequence homologous to the *Jumonji 14 (JMJ14)* gene. This suggested that the marker mutation itself might be causing earliness, as *AtJMJ14* is a negative regulator of flowering (Jeong et al., 2009; Yang et al., 2010; Lu et al., 2010). We annotated the *TaJMJ14* gene structure using the transcriptome data below (Figure 1a, Supplemental Figure 1a) and predicted the coding sequence (CDS) using the FGNESH gene structure prediction tool (Solovyev et al., 2006). Using this annotation, the earliness marker was a +C insertion 25 bp upstream of the *TaJMJ14* translational start site in the predicted 5’ untranslated region (UTR) (Figure 1a, Supplemental File 1).

**Figure 1.**
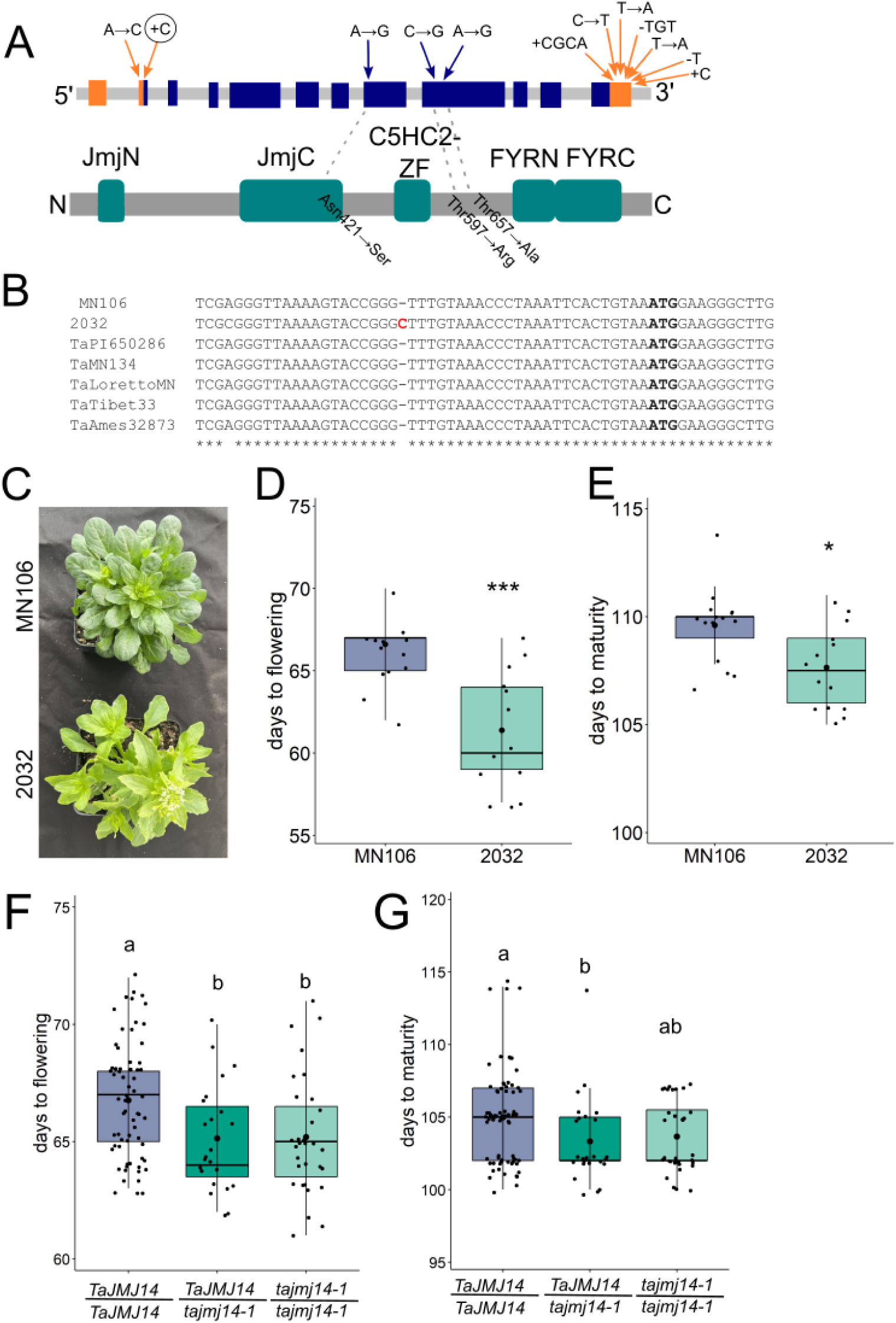
Early flowering and maturity phenotypes of 2032 plants co-segregate with mutations in the *TaJMJ14* gene. **(A)** Top: Diagram of the *TaJMJ14* gene. Larger rectangles represent exons; in orange are the 5’ and 3’ UTRs, and in blue is the predicted CDS. Bottom: Functional domains in the TaJMJ14 protein predicted by InterPro. Arrows indicate the location and type of mutations in the 2032 *TaJMJ14* gene compared to the reference MN106 sequence. Orange arrows highlight the location of UTR mutations. Circled is the +C insertion earliness marker. Gray dashed lines connect non-synonymous SNPs (indicated with blue arrows) to their corresponding amino acid substitutions, which are overlayed onto the protein model. Other intronic, promoter, and non-synonymous CDS mutations can be found in Supplemental File 1. All of the mutations in the 2032 *TaJMJ14* gene are referred to as the *tajmj14-1* allele. **(B)** Sequence alignment of part of the second exon of *TaJMJ14,* showing the earliness marker in red and the predicted start codon in bold. **(C)** Representative images of MN106 and 2032 plants during the flowering window (62 days after planting, including a 21-day vernalization period). Days from planting to when the first flower opened **(D)** and that the first dry, mature pod appeared **(E)**. Error bars = SD. * indicates a p<0.05 and *** indicate p<0.001 from a Student’s t-test. **(F-G)** 2032 and MN106 plants were crossed together, and 192 individual F2 plants were genotyped for the presence of the *tajmj14-1* allele and monitored for the appearance of the first flower and of the first mature pod. Bars = SD; groups indicated by the same letter are not significantly different as assessed by Kruskal-Wallis with Dunn’s post-hoc tests.

2032 is a natural pennycress accession with the earliness marker. When comparing the *TaJMJ14* gene sequence of other accessions in the pennycress pangenome (https://phytozome-next.jgi.doe.gov/pennypan/), the 5’UTR +C insertion associated with earliness is unique to 2032 (Figure 1b). Whole genome resequencing of 2032 found a variety of other mutations in the *TaJMJ14* gene compared to the reference MN106 sequence (Figure 1a), which are present in the pangenomes (Supplemental File 1). There was another 5’UTR mutation 43 bp upstream of the translational start site (Figure 1a,b). There were 3 non-synonymous substitutions in the predicted protein sequence, one of which, N421S, was in the conserved JmjC domain. There were 3 SNPs and 4 indels in the predicted 3’UTR (Figure 1a), as well as a variety of indels and SNPs in intronic and promoter regions (Supplemental File 1). All of the mutations in the 2032 *TaJMJ14* gene are henceforward referred to as the *tajmj14-1* allele.

We confirmed that the *tajmj14-1* allele conferred earliness in greenhouse studies by comparing the maturity of MN106 and 2032 plants. MN106 plants had their first open flower an average of 66.6 ± 3.0 days after planting, compared to 61.3 ± 3.6 days for 2032 plants (Figure 1c-d). In terms of maturity, 2032 plants had their first dry pod 107.8 ± 2.0 days after planting compared to 109.6 ± 1.8 days for MN106 plants (Figure 1e). Thus, in greenhouse conditions, 2032 plants with the *tajmj14-1* allele flowered on average 5 days earlier than MN106 plants leading to an earlier maturation of 2 days.

Next, we assessed the segregation of flowering time in 2032 x MN106 F2 plants. F2 plants homozygous for the reference (WT, MN106) *TaJMJ14* allele flowered on average 66.8 ± 2.5 days after germination. But those homozygous for the *tajmj14-1* allele flowered 1.5 days earlier, at an average of 65.2 ± 2.5 days after germination (Figure 1f). Homozygous *tajmj14-1* plants matured 1 day earlier than plants homozygous for the wild-type (WT) *TaJMJ14* allele; however, this difference was not statistically significant (Figure 1g). In an independent cross of 2032 with another CoverCress variety with a WT *TaJMJ14* allele, B36, homozygous *tajmj14-1* F2 plants flowered 5.8 days and matured 3.6 days earlier than homozygous *TaJMJ14-1* siblings, and both effects were statistically significant (Supplemental Figure 1b-c). Taken together, the co-segregation of the *tajmj14-1* allele with early flowering and maturity strongly suggests that TaJMJ14 is a repressor of flowering, similar to Arabidopsis JMJ14 (Jeong et al., 2009; Yang et al., 2010; Lu et al., 2010), and that *tajmj14-1* is a loss-of-function allele.

### The tajmj14-1 allele co-segregates with fungal disease susceptibility

In Arabidopsis, *jmj14* loss-of-function mutants are more susceptible to the bacterial pathogen *P. syringae* DC3000 (Li et al., 2020). Serendipitously, at the time the *tajmj14-1* allele was identified, various breeding populations derived from 2032 were being grown in the greenhouse and became infected with an unknown disease. It seemed that susceptibility to this disease was segregating in a Mendelian fashion. Plants from these populations were scored on a 1-5 scale for their health and genotyped for the presence of the *tajmj14-1* allele. Plants homozygous for the reference *TaJMJ14* allele were more likely to be lightly infected (score = 4) compared to plants homozygous for the 2032 *tajmj14-1* allele, which were more likely to be severely infected (score =2) (Figure 2a).

**Figure 2.**
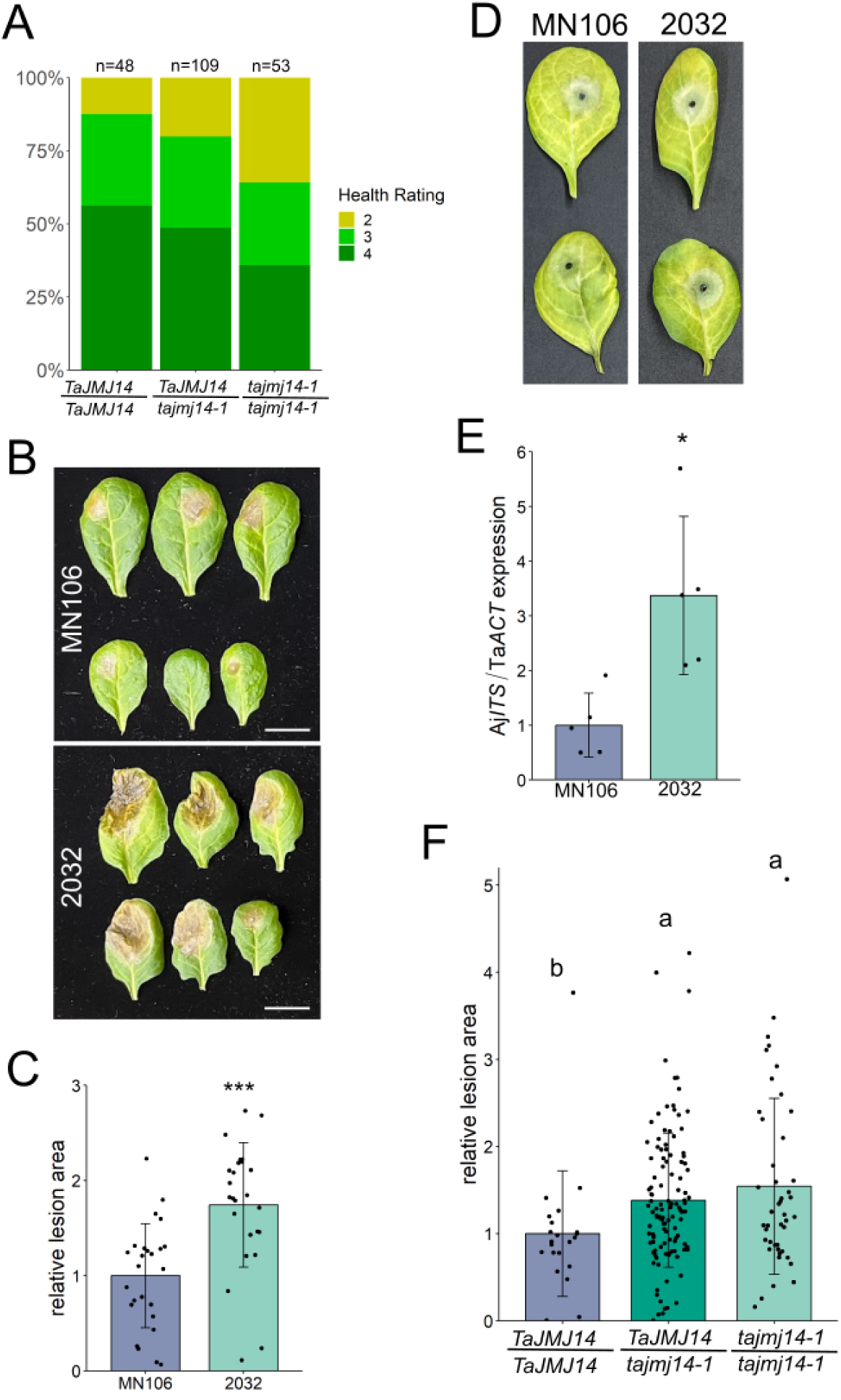
2032 leaves are more susceptible to *S. sclerotiorum* and *A. japonica* than MN106 leaves due to the *tajmj14-1* allele. **(A)** Segregating breeding populations derived from 2032 were scored for health using the following scale: 5=healthy, 4=light infection, 3=moderate infection, 2=severe infection, 1=dead. The same plants were genotyped for the presence of the *tajmj14-1* allele. **(B)** The 8th, 9th, and 10th leaves of 4.5 week-old non-vernalized plants were detached and inoculated with 2 mm agar plugs of *S. sclerotiorum*. Pictures of leaves from two plants per genotype were taken 3 dpi. Scale = 2 cm. **(C)** Quantification of the necrotic lesion area at 3 dpi relative to the mean lesion area on MN106 leaves. Each data point represents the mean lesion area of three leaves from one plant. Presented are the pooled results of three separate experiments. Bars = SD; *** represents p-value < 0.001 from Student’s t-test. **(D-E)** The 10^th^ leaves of plants were inoculated with 2 mm agar plugs of *A. japonica.* **(D)** Picture taken at 4 dpi. **(E)** *A. japonica* biomass on leaves 4 dpi as measured by qPCR detection of *A. japonica* DNA (Aj*ITS* gene) relative to pennycress DNA (Ta*ACT* gene). Expression was normalized by the -2^ΔΔCt^ method. Presented are the pooled results of two separate experiments, where each data point represents DNA from 3 pooled leaves. Bars = SD; * represents p-value < 0.05 from Student’s t-test. **(F)** 2032 and MN106 plants were crossed together, and 184 F2 plants were genotyped for the presence of the *tajmj14-1* allele. The 7^th^, 8^th^, and 9^th^ leaves of F2 plants were inoculated with 2 mm agar plugs of *S. sclerotiorum*. Plotted are the necrotic lesion areas (2-4 dpi) of individual F2 plants with each genotype (mean of 3 leaves), normalized to the mean lesion area of plants homozygous for the WT (MN106) *TaJMJ14* allele. Pooled are the results of 8 experiments. Bars = SD; groups indicated by the same letter are not significantly different as assessed by Kruskal-Wallis with Dunn’s post-hoc tests.

This led us to develop systematic, quantitative assays to test the effect of the *tajmj14-1* allele on disease resistance. We focused on two fungal pathogens, *S. sclerotiorum* and *A. japonica,* isolated from CoverCress fields (Kujur and Codjoe et al., 2025). Leaves of non-vernalized MN106 and 2032 plants were detached and inoculated with mycelial plugs of a field isolate of *S. sclerotiorum* and the brown, necrotic lesions that developed were measured. The lesion area at 3 days post-inoculation (dpi) was significantly larger on 2032 leaves than MN106 leaves (Figure 2b-c), indicating that 2032 leaves are more susceptible to *S. sclerotiorum.* Likewise, pods and stems from 2032 plants were also more susceptible to *S. sclerotiorum* than those from MN106 plants (Kujur and Codjoe et al., 2025).

When detached MN106 and 2032 leaves and pods were inoculated with *A. japonica* mycelial plugs, it was difficult to reliably see differences in fungal growth with the naked eye (Figure 2d). Henceforth, we developed a molecular assay to quantify *A. japonica* DNA as a proxy for fungal biomass (Kujur and Codjoe et al., 2025). Using this assay, we detected 3.4 times more *A. japonica* DNA on 2032 leaves than MN106 leaves 4 dpi (Figure 2e). *A. japonica* biomass is also greater on pods of 2032 compared to MN106 (Kujur and Codjoe et al., 2025). Taken together, these results indicate that in a variety of tissues and developmental stages, 2032 plants are more susceptible to *S. sclerotiorum* and *A. japonica* than MN106 plants.

To confirm that it was the *tajmj14-1* allele in 2032 plants that was responsible for fungal susceptibility, we screened the resistance of F2 plants derived from a cross between 2032 and MN106 to *S. sclerotiorum*. F2 plants homozygous for the 2032 *tajmj14-1* allele had infected lesions 54% larger than those homozygous for the reference *TaJMJ14* allele. Heterozygotes had a susceptibility statistically indistinguishable from *tajmj14-1* homozygotes (Figure 2f). This co-segregation strongly suggests that the *tajmj14-1* allele confers susceptibility to *S. sclerotiorum.* In Arabidopsis, *jmj14* loss-of-function mutants have impaired immunity to the bacterial pathogen *Pseudomonas syringae* DC3000 (Li et al. 2020), again suggesting that *tajmj14-1* is a loss-of-function allele.

### An independent CRISPR-induced allele confirms that disruption of the TaJMJ14 gene confers S. sclerotiorum resistance and possibly early flowering

We sought additional evidence that TaJMJ14 regulates flowering time and disease resistance in pennycress by generating new loss-of-function alleles in the gene using CRISPR-Cas9 gene editing technology. Gene editing was performed in a CoverCress line, B28 WG, with *tt8* and *fae1* knockout edits (McGinn et al., 2019; Sedbrook and Durrett, 2020; Gautam et al., 2025) using two guide RNAs targeting *TaJMJ14.* Two alleles created from the second protospacer were recovered in the T2 generation. One allele was a 9 bp deletion in the predicted 6^th^ exon of *TaJMJ14* (Figure 3a) which likely causes partial loss-of-function to the TaJMJ14 protein. We will refer to this line as *tajmj14-2.* The other allele, *tajmj14-3,* likely causes total loss-of-function, as it is a 16 bp deletion in the 6^th^ exon predicted to cause early termination before translation of the JmjC functional domain. A T2 plant homozygous for the *tajmj14-3* allele produced small, misshapen seeds, some of which appeared to have germinated before desiccation (Figure 3b). Upon rehydration, none of the seeds germinated, thus further characterization of *tajmj14-3* plants was not possible. However, the homozygous *tajmj14-2* T2 plant was fully fertile and made normal seeds (Figure 3b). T3 *tajmj14-2* plants were otherwise morphologically indistinguishable from parental control plants (Supplemental Figure 2).

**Figure 3.**
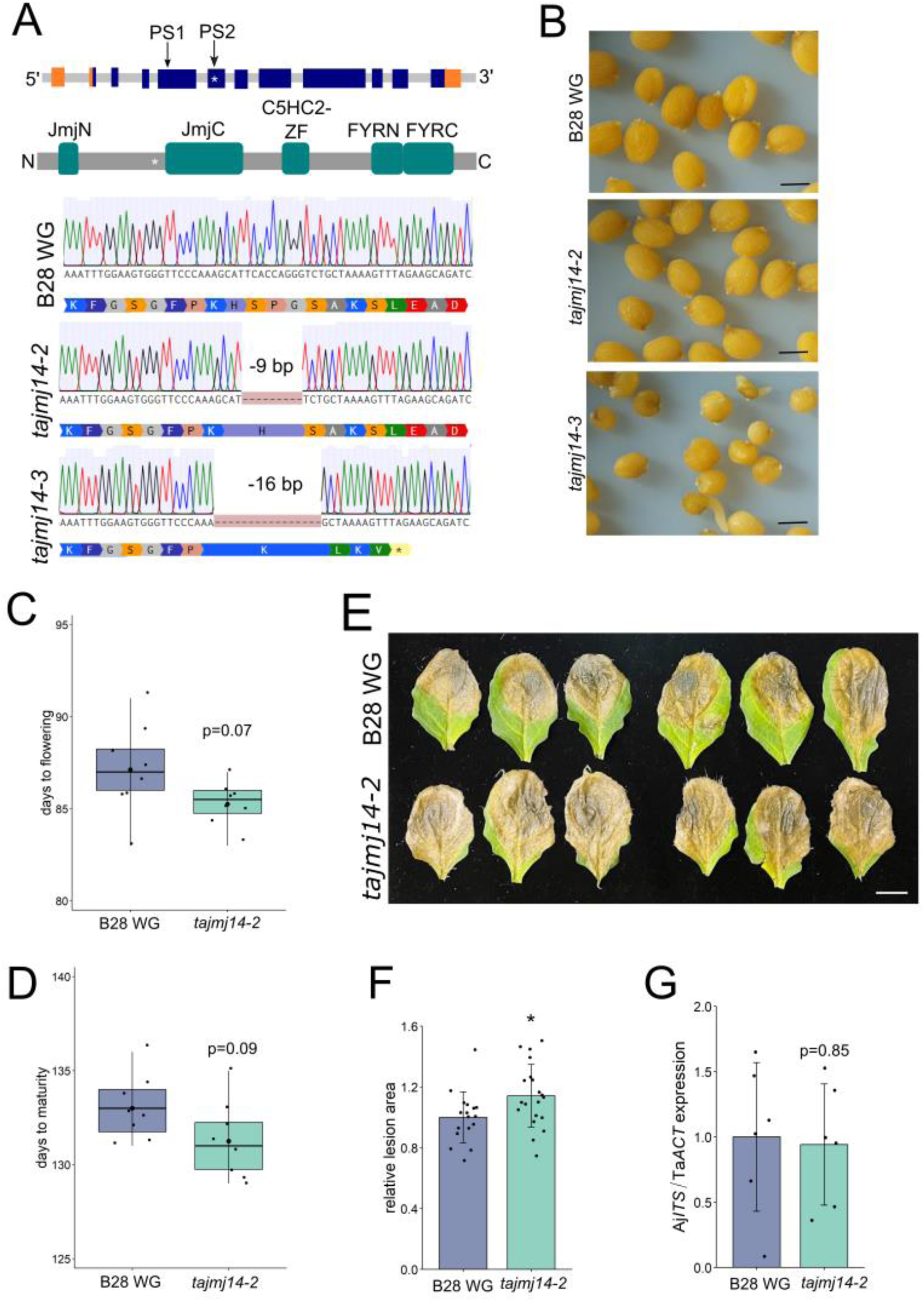
An independent, CRISPR-induced mutation in *TaJMJ14* confers susceptibility to *S. sclerotiorum* and potentially an early flowering phenotype. **(A)** Top: Diagram of the *TaJMJ14* gene and protein, as in Figure 1. Arrows indicate the location targeted by two protospacers (PS), and the asterisks indicate the location of the CRISPR-induced deletions in *tajmj14-2* and *tajmj14-3* plants. Bottom: Chromatograms and predicted translation of a region of the 6^th^ exon of *TaJMJ14,* showing a 9 bp deletion in *tajmj14-2* plants and a 16 bp deletion in *tajmj14-3* plants compared to parental (B28 WG) plants. **(B)** T3 seeds from *tajmj14-2* and *tajmj14-3* lines and a co-grown B28 WG plant. **(C-D)** Days from planting to the appearance of the 1^st^ flower (days to flowering) and the 1^st^ dry pod (days to maturity). Bars = SD; p-values from Student’s t-test. **(E-F)** The 7^th^, 8^th^, and 9^th^ leaves of 5 week old plants were inoculated with 2 mm agar plugs of *S. sclerotiorum*. **(E)** Pictures of leaves from two plants per genotype taken at 3 dpi. Scale = 2 cm. **(F)** Quantification of the necrotic lesion area at 3 dpi relative to the mean lesion area on parental B28 WG leaves. Each data point represents the mean lesion area of three leaves from one plant. Presented are the pooled results of two separate experiments. (G) *A. japonica* biomass as measured by qPCR detection of *A. japonica* DNA (Aj*ITS* gene) relative to pennycress DNA (Ta*ACT* gene). The 10^th^ leaves of plants were inoculated with 2 mm agar plugs of *A. japonica*, and infected leaf tissue was collected 4 dpi for DNA isolation. Expression was normalized by the -2^ΔΔCt^ method. Presented are the pooled results of two separate experiments, where each data point represents DNA from 3 pooled leaves. **(F-G)** Bars = SD; * represents p-value < 0.05 from Student’s t-test.

We tested the earliness and disease susceptibility of *tajmj14-2* plants. In growth chambers which mimicked the gradual increase in day length and temperature that pennycress plants in the field would experience, *tajmj14-2* plants flowered and matured on average 1.8 days earlier than B28 WG control plants, although not significantly so (Figure 3c-d). Three days after *S. sclerotiorum* inoculation, *tajmj14-2* leaves had 14% larger necrotic lesions than control leaves (Figure 5e-f), indicating that they had increased susceptibility to the pathogen. However, there was equivalent *A. japonica* biomass on *tajmj14-2* and control leaves at 4 dpi (Figure 3g). Although the plants with the *tajmj14-2* allele do not have all of the same phenotypes as *tajmj14-1* plants (i.e *A. japonica* susceptibility and a non-significant reduction in flowering time), our observation that two independent loss-of-function *tajmj14* alleles affect flowering time, maturity, and resistance to fungal pathogens supports the above evidence that these phenotypes are controlled by *TaJMJ14*.

### RNAseq identifies differentially expressed genes in 2032 and MN106 plants infected with A. japonica and S. sclerotiorum

As a histone demethylase, TaJMJ14 likely influences the expression of many genes, some of which promote disease resistance. We investigated the transcriptomes of MN106 and 2032 to identify potential resistance and susceptibility factors that may explain the differences in their susceptibility to fungal diseases. For RNAseq, we inoculated plants with either *S. sclerotiorum* or *A. japonica* to identify genes differentially expressed by both pathogens to narrow down those that might be most important for broad-spectrum fungal resistance and susceptibility. We focused our search on transcriptional changes occurring before disease establishment, i.e. prior to the onset of necrosis. Pennycress leaves first exhibited necrosis 12 hr post-inoculation (hpi) with *S. sclerotiorum* (Supplemental Figure 3a-b), as has been observed in *Brassica napus* (Novakova et al 2014). For *A. japonica,* we first observed necrosis at 24 hpi. Thus, we chose time points of 6 and 12 hpi for *A. japonica* and 3 and 6 hpi for *S. sclerotiorum* for RNAseq.

Fungal infection of pennycress led to more differential gene expression as the infections progressed. To illustrate, at 3 hr of *S. sclerotiorum* infection there were no differentially expressed genes (DEGs) in infected versus control MN106 plants, but by 6 hpi, there were 2810 DEGs (Figure 4a). At the same time point (6 hpi), differential gene expression is lower in *A. japonica-*infected leaves (MN106 has 261 DEGs) compared to *S. sclerotiorum-*infected leaves (MN106 has 2810 DEGs), consistent with the slower development of necrotic symptoms by *A. japonica*. Strikingly, in all cases, 2032 plants have more DEGs upon infection than MN106 plants. For example, at 12 hpi by *A. japonica* there are 6276 DEGs in 2032 leaves compared to 3232 DEGs in MN106. For all time points and for both pathogens, fewer genes are downregulated by infection than upregulated. Most of the genes induced by infection are common to both pathogens and both genotypes, whereas the downregulated genes are more unique to each pathogen and genotype than shared (Figure 4b).

**Figure 4.**
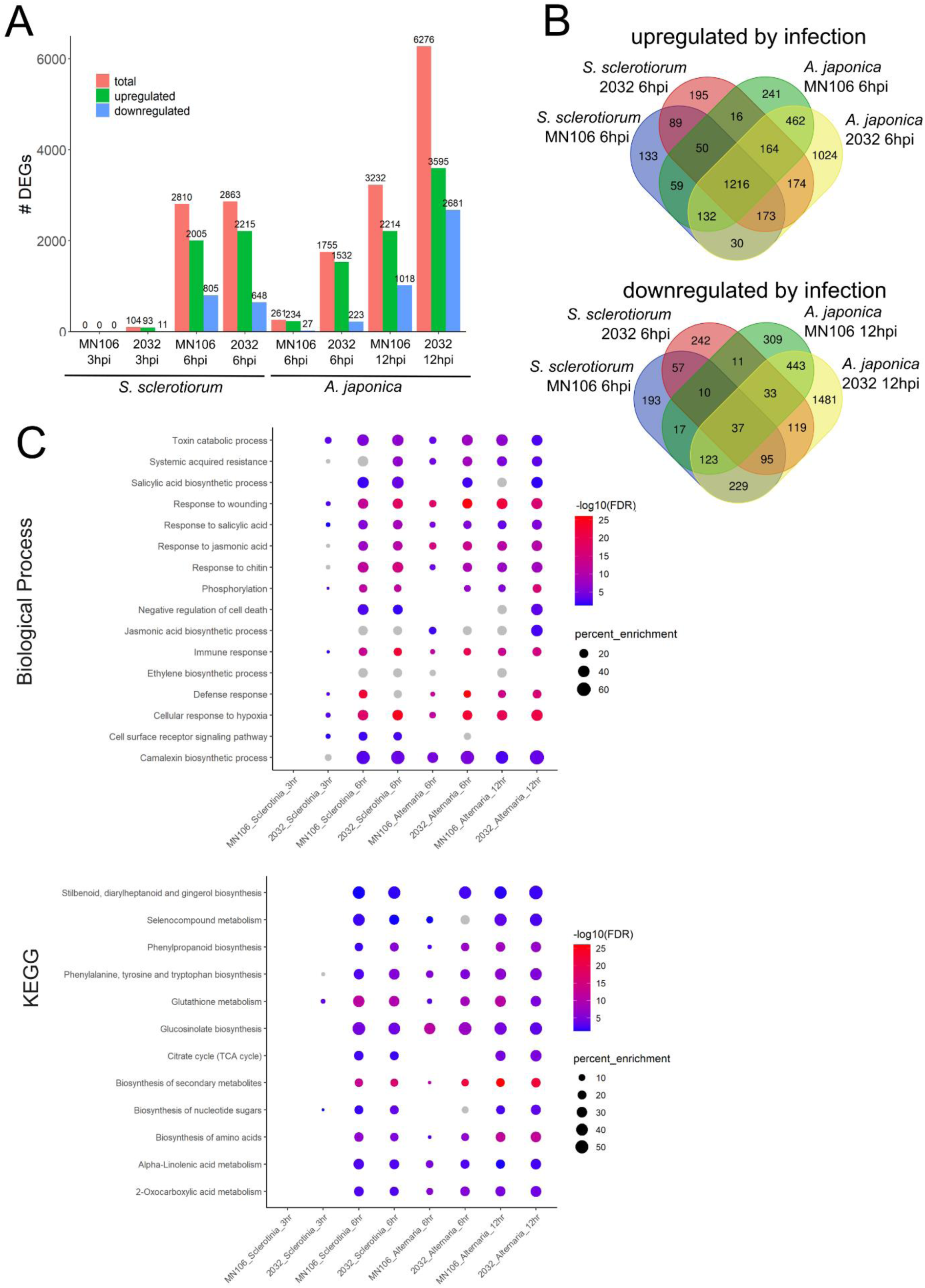
Greater transcriptional changes occur in infected 2032 compared to infected MN106 leaves, although both genotypes mount a fungal defense response. The 8^th^ rosette leaf of 5-week old MN106 and 2032 plants were infected with *S. sclerotiorum* (for 3 or 6 hr), *A. japonica* (for 6 or 12 hr), or sterile agar plugs for each time point. Total RNA was isolated from leaf tissue under and immediately surrounding the agar plugs and subjected to RNAseq. **(A)** Number of differentially expressed genes (DEGs) in each infection as compared to its mock control infection of the same time point. **(B)** The shared and unique DEGs for each infection. **(C)** The DEGs upregulated in each infection were subjected to gene ontology analysis. Presented are selected biological processes related to defense and the most enriched KEGG pathways. Dots are grey if the p-value cutoff (FDR) is > 0.05 and are absent if that pathway is not enriched in that infection.

In general, gene ontology (GO) analysis suggests that MN106 and 2032 plants share a similar response to fungal infection. Biological process (BP) GO terms related to fungal defense are significantly enriched in both infected MN106 and infected 2032 samples. These GO terms include defense response, immune response, response to wounding, and the responses to salicylic acid (SA) and jasmonic acid (JA), key hormones in defense against *S. sclerotiorum* (Guo et al., 2007) and *Alternaria* species (Brouwer et al., 2020; Nair et al., 2014; Figure 4c). The number of genes represented from each pathway differs slightly between genotypes, but in general both MN106 and 2032 have enrichment of similar pathways. Likewise, similar KEGG pathways are enriched in the induced DEGs of both genotypes, including those involved in glucosinolate and phenylpropanoid biosynthesis and 2-oxocarboxylic acid, α-linolenic acid, and glutathione metabolism. Increased glucosinolate and phenylpropanoid biosynthesis in response to *S. sclerotiorum* infection contributes to resistance in *B. napus* and soybean, respectively (Wu et al., 2016; Ranjan et al., 2019). α-linolenic acid metabolism contributes to the production of JA and volatile compounds (Chen et al., 2023), which may contribute to fungal resistance. The activity of some glutathione-S-transferases can promote resistance to *S. sclerotiorum* and other pathogens (Chen et al., 2021; Gullner et al., 2018). In both MN106 and 2032 leaves, the types of pathways downregulated by infection are related to photosynthesis, development, and primary metabolism (Supplemental Figure 3c).

### RNAseq reveals differences in gene expression between 2032 and MN106 plants that may underlie their differing susceptibilities to fungal diseases

Despite the apparent activation of disease response pathways in infected 2032 leaves, 2032 plants are immunocompromised. In our investigation to understand why 2032 plants are more vulnerable to disease, we searched for susceptibility factors in the genes with significantly higher expression in infected 2032 compared to infected MN106 plants. Depending on the infection, between 1445 and 2308 genes are more significantly expressed in infected 2032 leaves than infected MN106 leaves (Supplemental Figure 4a). There was no enrichment of KEGG pathways or BP GO terms in these subsets of genes. To narrow down the list to those genes that might confer broad-spectrum susceptibility, we focused on the 249 genes that were higher in 2032 leaves during both *S. sclerotiorum* and *A. japonica* infection, (Supplemental File 2). Such genes may be direct targets of JMJ14, as the presumed loss-of-function of JMJ14 in 2032 plants should lead to H3K4me3 accumulation and transcriptional activation of target genes.

One of the potential susceptibility factors in 2032 plants is a gene with homology to *COMPASS-like H3K4 histone methylase component WDR5a* (TAV2_LOCUS12666, AT2G37670), whose mRNA expression was higher in 2032 than MN106 leaves both constitutively and under infection (Figure 5a). In Arabidopsis, WDR5a and WRKR5b trimethylate histone H3K4 (Jiang et al., 2009, 2011), the same histone mark that JMJ14 demethylates (Lu et al., 2010b; Yang et al., 2018). Higher expression of *WDR5a* in 2032 plants may exacerbate the effects of the loss of JMJ14 function. Wall-associated kinase-like protein 11 (WAKL11, TAV2_LOCUS3598, AT1G19390) promotes pectin degradation in Arabidopsis (G. H. Han et al., 2023). *S. sclerotiorum* and other pathogenic fungi produce pectin- and other cell wall-degrading enzymes as pathogenicity factors (Bolton et al., 2006). The higher *WAKL11* expression in 2032 plants may promote infection by further reducing pectin levels. Finally, serine/threonine/tyrosine kinase 17 (STY17, TAV2_LOCUS5673, AT4G35780), whose mRNA expression is constitutively higher in 2032 compared to MN106 plants, is a negative regulator of jasmonate and chitin responses and confers susceptibility to the fungal necrotroph *Botrytis cinerea* (Chen et al., 2024).

**Figure 5.**
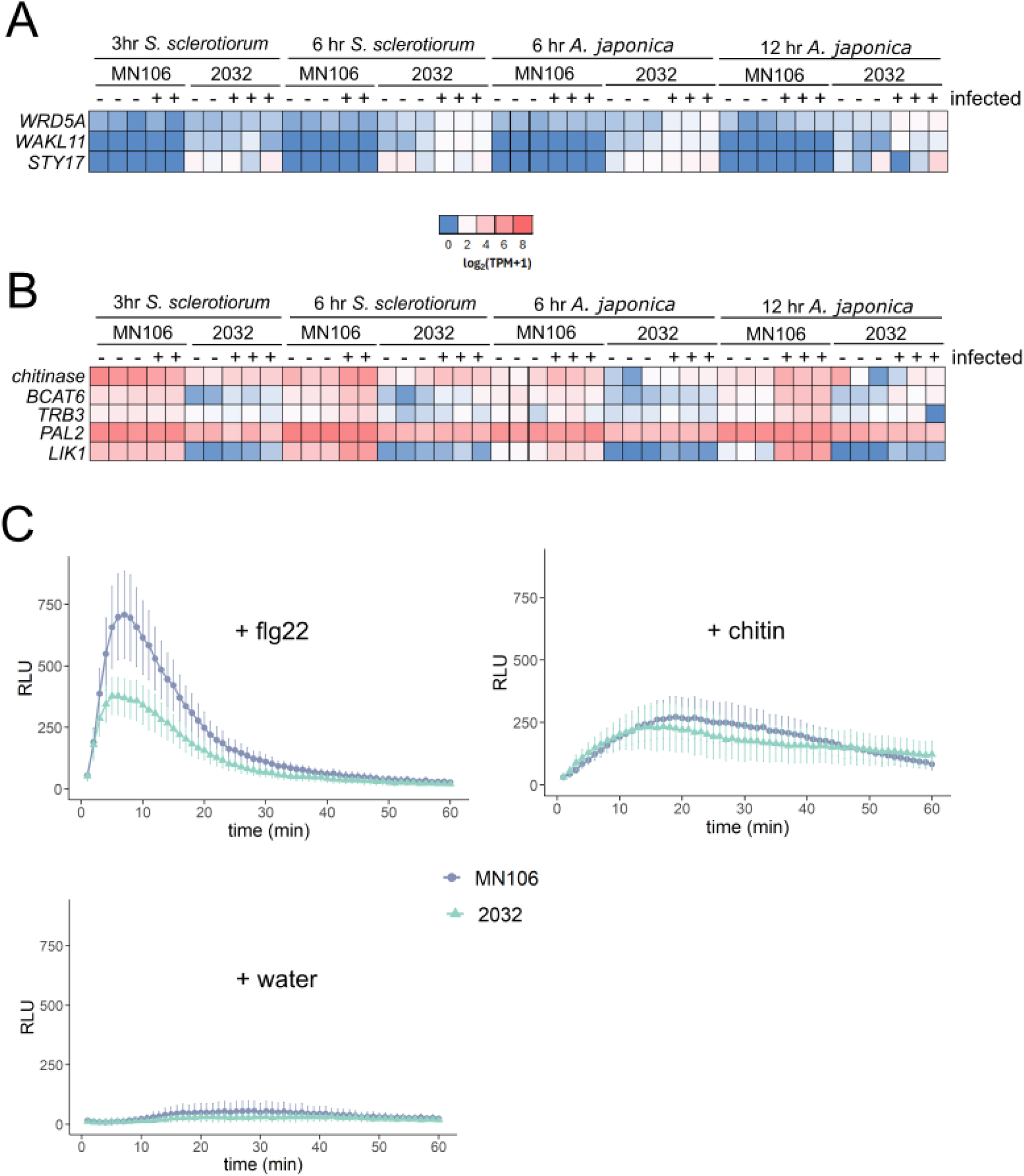
Compared to MN106, 2032 leaves have higher expression of putative susceptibility genes, lower expression of putative resistance genes, and exhibit weaker oxidative bursts in response to PTI elicitors. (A-B) Expression of selected genes displayed in log_2_(transcripts per million +1) form. Genes in **(A)** have significantly higher expression in infected 2032 than infected MN106 samples across all 4 infections (two pathogens, two time points) according to DESeq2 analysis. Similarly, genes in **(B)** have significantly higher expression in infected MN106 than infected 2032 samples for all infections. **(C)** Leaf discs were collected from the 6th leaves of 5-week-old MN106 and 2032 plants, incubated in a luminol-HRP solution, and treated with water,1 µM flg22, or 10 µg/mL chitin **(C).** Relative luminescence units (RLU) were recorded for 1 hr. Plotted are the mean RLU ± SEM from 4 separate experiments.

Conversely, to find the genes that might contribute to resistance in MN106 plants, we searched the list of 489 genes that were significantly upregulated in infected MN106 plants compared to infected 2032 plants across all infections (Supplemental File 2), looking for those that had reported or predicted roles in defense. These include a putative chitinase (TAV2_LOCUS12783, AT2G43570), which may degrade fungal cell walls, and branched-chain-amino-acid aminotransferase 6 (BCAT6, TAV2_LOCUS2477, AT1G50110), which functions in the biosynthesis of methionine-derived aliphatic glucosinolates (Lachler et al., 2015) (Figure 5b). The breakdown products of aliphatic glucosinolates restrict the growth of *A. brassicicola* (Buxdorf et al., 2013) and *S. sclerotiorum* (Stotz et al., 2011; Chen et al., 2020). The higher expression of *telomere repeat-binding factor 3* (*TRB3*, TAV2_LOCUS15987, AT3G49850) in MN106 plants may enhance JMJ14-mediated immunity as it is known to recruit JMJ14 and PRC2 to target loci to repress gene expression (Wang et al., 2023). Phenylalanine ammonia lyase 2 (PAL2, TAV2_LOCUS25534, AT3G53260) functions redundantly in phenylpropanoid, including salicylic acid (SA) biosynthesis (Huang et al., 2010). The higher *PAL2* expression in infected MN106 leaves would be predicted to contribute to fungal defense as SA promotes resistance to *S. sclerotiorum* (Guo and Stotz, 2007) and *A. brassicae* (Joshi et al., 2010) and many metabolites and lignin derived from the phenylpropanoid pathway are associated with resistance to *S. sclerotiorum* (Eynck et al., 2012; Jain et al., 2015; Chen et al., 2019; Ranjan et al., 2019).

We note that there are several nucleotide-binding leucine-rich repeat (NLR) proteins among the genes most highly expressed in infected MN106 versus infected 2032 samples (Supplemental Figure 4a, Supplemental File 2), but it is extremely rare for R-genes to confer resistance to necrotrophic pathogens (Ghozlan et al., 2020), and it is unlikely that any are contributing to resistance here in MN106 plants. Arabidopsis *jmj14* mutants are more susceptible to the hemibiotrophic bacterial pathogen *P. syringae,* perhaps in part due to compromised activation of defense-related gene expression during infection (Li et al., 2020). We observed a similar pattern as reported in Li et al. (2020) in which the expression of *pathogenesis-related 1 (PR1),* a marker activated by salicylic acid signaling (Z. Han et al., 2023), and *AGD2-like defense response protein 1 (ALD1),* a pipecolic acid biosynthesis gene (Návarová et al., 2012), is induced by infection and observed that this induction is attenuated in *tajmj14-1* (2032) plants (Supplemental Figure 4b). We did not see this effect for all *jmj14-*dependent genes reported in Li et al. (2020) in our experiments (Supplemental Figure 4b), perhaps because of the different pathogens or time points involved.

One of the most enriched genes in *A. japonica-*infected MN106 leaves at 12 hpi was *LysM-RLK1-interacting kinase 1* (*LIK1,* TAV2_LOCUS7511, AT3G14840) (Supplemental Figure 4a). *LIK1* is also expressed significantly higher in MN106 leaves in all infections (Figure 5b, Supplemental File 2). LIK1 functions in cell-surface perception of fungal chitin and bacterial flagellin in Arabidopsis, activating signaling that confers resistance to *S. sclerotiorum* (Le et al., 2014). LIK1 interacts with chitin elicitor receptor kinase 1 (CERK1) (Le et al., 2014), which promotes immunity to *A. brassicicola* (Miya et al., 2007)*. LIK1* expression is induced by infection in both genotypes but to a higher level in MN106, and its expression is constitutively higher in MN106 compared to 2032 samples (Figure 5b). This led us to test if PAMP-triggered immunity was compromised in 2032 leaves. Indeed, the oxidative bursts triggered by the flg22 peptide, and to a lesser extent, chitin, were reduced in 2032 leaves (Figure 5c).

## DISCUSSION

The genetic and phenotypic diversity found in natural accessions represents a valuable resource for plant breeding. Breeders can identify naturally occurring alleles associated with desirable agronomic traits to improve yield, stress resilience, and adaptation to specific local environments. 2032, a natural accession of *T. arvense* isolated in Colorado, USA, has been a valuable resource for CoverCress breeding as it has considerable genomic divergence from other North American pennycress accessions. One of the valuable agronomic traits associated with 2032 is early flowering, and to a lesser extent, early maturity (Figure 1). Pennycress matures and is harvested in the late spring in the U.S. Midwest, a time when farmers are otherwise planting corn or soybeans. Earlier maturing pennycress varieties minimize potential disruptions to the growing season of these subsequent crops.

In greenhouse conditions, 2032 plants flower 5 days and mature 2 days earlier than another North American pennycress accession, MN106. We demonstrated that at least some of that earliness comes from naturally occurring mutations in the *TaJMJ14* gene in 2032 plants, which we termed the *tajmj14-1* allele (Figure 1). In a segregating 2032 x MN106 F2 population, the *tajmj14-1* allele conferred an earlier flowering phenotype of only 1.5 days, suggesting that there are other elements in the 2032 genome that contribute to its early flowering phenotype. However, the effect of the *tajmj14-1* allele may be complicated by epistatic interactions in that cross, as the *tajmj14-1* allele conferred an early flowering phenotype of 5.8 days in a cross of 2032 to a CoverCress variety, B36 (Supplemental Figure 1). Previous studies that implicated JMJ14 in repressing the floral transition were done in the Col-0 ecotype of Arabidopsis (Jeong et al., 2009; Lu et al., 2010; Yang et al., 2010), which exhibits summer annual behavior. To our knowledge, this study is the first to show that JMJ14 also delays the flowering transition in a species that requires vernalization for flowering.

Yet, the early flowering and maturity associated with the *tajmj14-1* allele may not be useful for pennycress breeders, as the allele also causes susceptibility to several diseases (Figure 2). We first discovered this phenomenon accidentally; plants with the *tajmj14-1* allele in breeding populations derived from 2032 were more vulnerable to an unknown disease in the greenhouse compared to their siblings with the WT *TaJMJ14* allele. We then confirmed its effect on disease susceptibility in a controlled experiment using the fungal pathogen *S. sclerotiorum,* the causal agent of white mold disease. The severity of *S. sclerotiorum* disease symptoms on leaves co-segregates with the *tajmj14-1* allele in F2 plants from a 2032 x MN106 cross (Figure 2). Presumably it is the *tajmj14-1* allele that is responsible for the increased susceptibility to *S. sclerotiorum* that we also observed on 2032 pods and stems as well as the susceptibility of 2032 leaves and pods to *A. japonica* (Figure 2; Kujur and Codjoe et al., 2025), although this would need to be confirmed. *JMJ14* is ubiquitously expressed in Arabidopsis (BAR eFP Browser, (Winter et al., 2007)).

2032 has both silent and non-synonymous mutations in the predicted CDS of the *TaJMJ14* gene, as well as non-coding mutations in the 5’ and 3’UTRs, introns, and promoter regions (Figure 1a, Supplemental File 1). Because these mutations are presumably linked, we refer to them as a whole as the *tajmj14-1* allele. Considering that Arabidopsis null *jmj14* mutants flower early and have impaired disease resistance (Jeong et al., 2009; Lu et al., 2010; Yang et al., 2010; Li et al., 2020), we infer that *tajmj14-1* is a loss-of-function allele. However, we do not yet know which mutation, or which combination of mutations, in the *tajmj14-1* allele is causal. One SNP causes a N421S substitution in the predicted JmjC domain. The JmjN, JmjC, and C5HC2 domains together function in substrate recognition and contain the catalytic residues for H3K4me3 demethylase activity (Yang et al., 2018). Alignment with Arabidopsis JMJ14 suggest that N421 in TaJMJ14 is not one of the catalytic residues, but any of the 3 non-synonymous substitutions in 2032 could potentially impair TaJMJ14 demethylase activity or its recruitment to target loci by interactions with NAC transcription factors (Zhang et al., 2015) or TRB proteins (Wang et al., 2023). The earliness marker, a single base pair insertion in the *TaJMJ14* 5’UTR, might be causal, as it is unique to 2032 in the pennycress pangenome (Figure 1b, Supplemental File 1), and CoverCress Inc. has found that 2032 plants flower earlier in the field than any of the accessions in the pangenome. In our RNAseq experiments, the expression of the *TaJMJ14* transcript is not significantly different in MN106 and 2032 leaves (Supplemental Figure 5), suggesting that the non-coding mutations have no effect on *TaJMJ14* transcription or mRNA stability. However, the mutations in the 5’UTR and/or 3’UTR could be reducing *TaJMJ14* translation.

Our generation of another presumed loss-of-function mutant, *tajmj14-2,* using gene editing provided additional evidence that pennycress JMJ14 functions to positively regulate disease resistance and negatively regulate flowering time (Figure 3). The *tajmj14-2* allele is a 3 amino acid deletion just outside of the JmjC domain in a region of the protein that is considered part of the overall Jumonji catalytic domain (Yang et al., 2018). The *tajmj14-2* allele appears to be weaker than the *tajmj14-1* allele, as *tajmj14-2* plants were only slightly more susceptible to *S. sclerotiorum* infection, and not at all to *A. japonica*. Additionally, *tajmj14-2* plants flower and mature earlier than parental control plants, but not significantly so. Overall, this suggests that *tajmj14-2* is a partial loss-of-function allele. We were unable to characterize full *TaJMJ14* knockout plants due to infertility (*tajmj14-3;* Figure 3a-b). The sterility associated with the presumed loss of TaJMJ14 protein function in the *tajmj14-3* line suggests that the *tajmj14-1* allele causes partial loss-of-function to TaJMJ14, as *tajmj14-1* plants produce fertile seeds.

RNAseq allowed a deeper understanding of how reduced TaJMJ14 function leads to fungal disease susceptibility. Even without proper epigenetic reprogramming by TaJMJ14, 2032 plants infected with *S. sclerotiorum* and *A. japonica* still mount a strong transcriptional response against pathogens, at least at the level of pathway analysis (Figure 4c). Additionally, for each infection there were no GO pathways enriched 2032 versus MN106 samples, or vice versa, to implicate any pathway as contributing to the differences we observed in fungal immunity. Similarly in *B. napus*, similar sets of defense-related genes are induced by *S. sclerotiorum* in both resistant and susceptible cultivars; the differences in susceptibility are attributed more to the expression level of genes in those pathways (Wu et al., 2016).

Fungal infection appeared to proceed more rapidly in 2032 plants, as there were consistently more DEGs in infected 2032 compared to infected MN106 leaves at each time point (Figure 4a). This is not surprising considering that 2032 leaves had dampened PTI, in terms of elicitor-triggered oxidative bursts (Figure 5c). PTI signaling includes, but is not limited to, the activation of SA and jasmonate signaling, secondary metabolite production, and cell wall remodeling to inhibit pathogen growth (Bigeard et al., 2015; Lloyd et al., 2014). Some of the potential susceptibility and resistance factors explaining the immunity differences in MN106 and 2032 to the two fungal pathogens can be attributed to genes functioning in these downstream PTI categories. Those highlighted in Figure 5 would be leading candidates for gene editing to improve disease resistance in pennycress. Knocking out the putative susceptibility genes and/or boosting the expression of the putative resistance factors in 2032 plants has the potential to restore a wild-type level of resistance to fungal pathogens while keeping intact the favorable early flowering phenotype.

Here we characterized just two traits associated with TaJMJ14, but there are likely many more considering its predicted function as a histone remodeler. While the pleiotropic effects of epigenetic regulators present challenges, as with disease susceptibility here, they also offer unique avenues for improving multiple agronomic traits simultaneously. By leveraging advanced genetic and epigenetic tools, future breeding efforts could separate the favorable traits from the negative consequences of such pleiotropy, ultimately improving the performance of pennycress in diverse agricultural systems.

## MATERIALS AND METHODS

### Plant Growth Conditions

Non-vernalized plants for disease assays were grown in growth chambers maintaining temperatures of 25°C in the day and 23°C at night, 300 µmol light intensity, 50% relative humidity, and a 14 hr day length. Seeds were soaked in 3.3 µg/mL gibberellic acid 4+7 overnight then sown in Pro-Mix FPX with Biofungicide germination mix. After approximately 2 weeks, seedlings were transplanted into pots containing Berger BM7 35% Bark mix.

For flowering studies, 2-week-old seedlings were vernalized at 4°C for 21 days, transplanted into pots containing Berger BM7 35% Bark mix, and grown in greenhouses with supplemental lights at 1000 µmol intensity, a 22 hr day length, and temperatures between 20 and 24°C. The exception was the flowering and maturity study in Figure 3c-d, which was designed to mimic some of the daylength and temperature changes pennycress plants experience in the field. Two-week old seedlings were vernalized for 4 weeks at 4°C in a growth chamber with a 10 hr day length. Subsequently the plants spent 2 weeks at 10°C with a 12 hr day length, 2 weeks at 15°C with 14 hr day length, and then 20°C in the day (16 hr) and 18°C at night (8 hr) until maturity.

### Fungal cultures

Isolation and identification of *S. sclerotiorum* and *A. japonica* strains from field grown pennycress is described in detail in Kujur and Codjoe et al., 2025. Every 6 weeks, a new *S. sclerotiorum* culture was started from sclerotia. A single sclerotium was sterilized in 50% bleach, cut open, and incubated on a water agar plate. Mycelia that grew from this sclerotium was transferred to a potato dextrose agar (PDA) plate and incubated at 25°C in the dark. This plate was used to begin subsequent cultures. *S. sclerotiorum* was subcultured by taking a punch (using the back of a sterile p200 pipette tip) of mycelia from one PDA plate to a fresh PDA plate. Fresh *A. japonica* mycelial cultures were started every 6 weeks from frozen stocks. Mycelia were spread on PDA plates supplemented with 0.02 mg/mL each of chloramphenicol and streptomycin sulfate, and plates were incubated at 25°C in the dark. Subcultures were prepared every two weeks by cutting a piece of infected PDA and transferring to fresh PDA plates with the above antibiotics.

### S. sclerotiorum infection assays

For detached leaf infection assays, rosette leaves were detached from 4.5- to 6-week-old non-vernalized plants and transferred to petri dishes lined with a paper towel and 2 mL sterile water. Mycelial agar plugs (2 mm in diameter) were taken from the leading edge of a plate of *S. sclerotiorum* 3 days after transfer to a fresh PDA plate and placed on the adaxial leaf surface, avoiding veins. Lids were placed on the petri dishes and sealed in a plastic storage tote lined with wet paper towels to maintain high humidity. Leaves were incubated at room temperature in the dark for up to 4 days. The area of necrotic lesions was quantified in FIJI software using images taken with a camera or by CropReporter (PhenoVation, Wageningen, Netherlands).

### A. japonica infection assay

The 10^th^ rosette leaves from 4.5- to 5-week old non-vernalized plants were detached and placed in petri dishes lined with wet paper towels. 2 mm mycelial agar plugs from the leading edge of a 14-day old *A. japonica* plate were placed on the adaxial leaf surface, avoiding veins. Lids were placed on the petri dishes and sealed in a plastic storage tote lined with wet paper towels to maintain high humidity, and incubated in the dark at room temperature. At 4 dpi, the agar plugs were removed, and 22 mm leaf discs of the infected tissue were collected and flash frozen. Genomic DNA was isolated using the Quick-DNA Plant/Seed Miniprep Kit (Zymo Research). The relative amounts of fungal and plant DNA in each sample were determined by quantitative qPCR, using primers to detect *A. japonica ITS* and *T. arvense actin* genes (Supplemental Table 1). 5 ng of genomic DNA was used as template for each reaction and amplified using the igSYBR Green qPCR 2X Master Mix (Intact Genomics).

### Genotyping tajmj14-1 allele in F2 and breeding populations

Genomic DNA was isolated from leaf tissue using the GenCatch Plant Genomic DNA Miniprep Kit (Epoch Life Science). The presence of the *tajmj14-1* allele in these plants was determined with a PCR Allelic Competitive Extension (PACE) assay using PACE 2.0 Genotyping Master Mix with low ROX (3CR Bioscience), an allele specific primer, a WT allele specific primer, and a common primer (Supplemental Table 1). These primers detect the presence or absence of the +C insertion in the *TaJMJ14* 5’UTR.

### Creation and genotyping of tajmj14-2 mutant

Two protospacers (5’-GAATTGTGCCGCCTGTTGCA-3’ and 5’-GGTTCCCAAAGCATTCACCA-3’) targeting *TaJMJ14* were cloned into a CoverCress vector containing gRNA scaffolds under the control of *AtU6-26* promoters and *Sp*Cas9 under the control of the *AtRPS5A* promoter, with a DsRed selectable marker. This construct was transformed into the B28 WG genotype of CoverCress using floral dip as previously described (McGinn et al., 2019). Transformed T1 seeds were identified by DsRed fluorescence. DsRed negative (T-DNA free) T2 seeds were genotyped using Sanger sequencing with primers in Supplemental Table 1. Flowering and fungal susceptibility experiments were performed on homozygous edited (*tajmj14-2*) T3 plants.

### S. sclerotiorum infection and RNA isolation for RNAseq

Five-week-old, non-vernalized MN106 and 2032 plants were inoculated with two 6 mm plugs from the leading edge of a 3 day old *S. sclerotiorum* culture (infected), or plugs from a sterile PDA plate (mock infected), on their 8^th^ rosette leaf. Infected and mock-infected plants were kept in a plastic tote lined with wet paper towels at room temperature. After 3 or 6 hours post-inoculation (hpi), 1 cm leaf punches were taken around the agar plugs. The agar plugs were removed and the leaf punches were flash frozen in liquid nitrogen. Leaf punches from 2 plants (4 punches in total) were pooled to make one biological replicate.

Leaf punches were homogenized in liquid nitrogen, and total RNA was isolated using the NucleoSpin Plant RNA Kit (Machery-Nagel) with lysis in Buffer RAP and an on-column DNase digestion, following the manufacturer’s instructions. RNA was concentrated using the precipitation protocol of Hughes (2019).

### A. japonica infection and RNA isolation for RNAseq

The 8^th^, 9^th^ and 10^th^ leaves of 4.5 week old non-vernalized MN106 and 2032 plants were inoculated with 2 or 3 plugs of *A. japonica* mycelia (6mm) from the growing edge of a 14-day-old subculture or a sterile PDA plate. Plants were then placed in airtight container lined with wet paper towels for 6 or 12 hpi. The plugs were removed and 8mm leaf discs were collected and flash frozen and stored at -80° C until processing. Each biological replicate sample contained a leaf disc from each of the 8^th^, 9^th^, and 10^th^ leaves from 2 plants (6 punches in total). Samples were homogenized in liquid nitrogen and total RNA extracted using the E.Z.N.A.® Plant RNA Kit (Omega Bio-tek, Inc.) by following the manufacturer’s instructions.

### RNA sequencing

mRNA enrichment, library preparation, and transcriptome sequencing were performed by Novogene. Sequencing libraries were generated from 1 ug of total RNA using the NEBNext Ultra RNA Library Prep Kit for Illumina (NEB) following manufacturer’s recommendations. Index codes were added to attribute sequences to each sample. The clustering of the index-coded samples was performed on an Illumina Novaseq 6000 sequencer according to the manufacturer’s instructions. After cluster generation, the libraries were sequenced on the same machine and 20 million 150 nt paired-end reads were generated per sample.

### Transcriptome analysis

Raw reads in FASTQ format were processed through fastp software (Chen et al., 2018). In this step, clean reads were obtained by removing those reads containing adapter, reads containing N > 10% or reads containing >50% low quality bases (Qphred ≤ 5). The *T. arvense* v2 reference genome and gene model annotation files were downloaded from NCBI (GenBank assembly GCA_911865555.2). An index of the reference genome was built and paired-end clean reads were aligned to it using Hisat2 v2.0.5 (Kim et al., 2019), a splice aware aligner. featureCounts v1.5.0-p3 (Liao et al., 2014) was used to count the reads mapped to each gene. The resulting gene count matrix and gene lengths were used to calculate transcripts per million (TPM) values using the DEGobj.utils R package. These TPM values were log transformed with a pseudo count of 1; log_2_(TPM + 1) values were used for principal components analysis (PCA). Three samples from the *Sclerotinia* dataset were dropped from subsequent analyses because considerable differences between other biological replicates as visualized on PCA plots.

Differential expression analysis of two groups was performed using the DESeq2 R package (v1.20.0; Love et al., 2014). The resulting p-values were adjusted using Benjamini and Hochberg’s approach for controlling the false discovery rate. Genes with an adjusted p-value ≤

0.05 and an absolute fold change ≥ 2 were assigned as differentially expressed. Venn diagrams of DEGs were constructed using https://bioinformatics.psb.ugent.be/webtools/Venn/. For gene enrichment analysis, the *T. arvense* transcripts were subjected to BLASTn searches against the *A. thaliana* TAIR10.1 genome and the top hit was recorded. These Arabidopsis orthologs of the pennycress differentially expressed genes (DEGs) were subjected to gene ontology (GO) analysis using ShinyGO v0.77 (Ge et al., 2020).

### Elicitor-triggered oxidative burst assays

4mm leaf punches were collected from the 6^th^ rosette leaves of 5-6-week-old plants, in technical duplicate, and placed in wells of a black 96-well plate containing 150 µL water per well and incubated in the dark overnight. The water was removed and replaced with 120 µL of 30 µg/mL luminol (Ward’s Science) and 20 µg/mL horseradish peroxidase (VWR) solution, and the plate was incubated in the dark for 30 min prior to the addition of elicitors (Trujillo, 2016). 120 µL of 2 µM flg22 (GenScript), 20 µg/mL chitin (from shrimp shells, Megazyme), or water was added to each well, and luminescence readings were recorded every minute for 60 minutes using a Tecan plate reader.

### Statistical analyses

Statistical analyses were performed in RStudio (v4.3.1). Shapiro–Wilk tests were used to test for normality. Normally distributed data was analyzed using Student’s t-test or ANOVA; non-normal data with the Kruskal-Wallis test. The car and agricolae packages were used to perform ANOVA and Scheffe’s post-hoc tests and the rstatix package for Dunn’s Multiple Comparison post-hoc test. Data was visualized using RStudio ggplot2 packages.

## ACCESSIONS

Raw and processed data from RNAseq was deposited to NCBI GEO with the series accession number GSE294781.

## AUTHOR CONTRIBUTIONS

D.M.S., T.U., and R.C. conceptualized and supervised the project. J.M.C., A.K., T.S., and A.S. collected data and performed the experiments. K.R. and J.M.C annotated genes, and K.R. designed editing constructs. J.P.S. and J.M.C performed bioinformatic analysis and RNAseq interpretation, with discussion and input from A.K. J.M.C. wrote the manuscript with editing input from all authors.

## ACKNOWLEDGEMENTS

Funding was provided by USDA-NIFA-SBIR Phase II grant 2021-06450. We would like the members of the CoverCress Transformation and Editing and QAQC Teams for transforming, screening, and revalidating the CRISPR-Cas9 edited lines. Thanks to Ray Kennedy from CoverCress for collecting some maturity data.

