## Supplementary Figures and Tables for "Fungal susceptibility and early flowering in pennycress (*Thlaspi arvense*) are conferred by naturally occurring mutations in histone demethylase Jumonji 14"

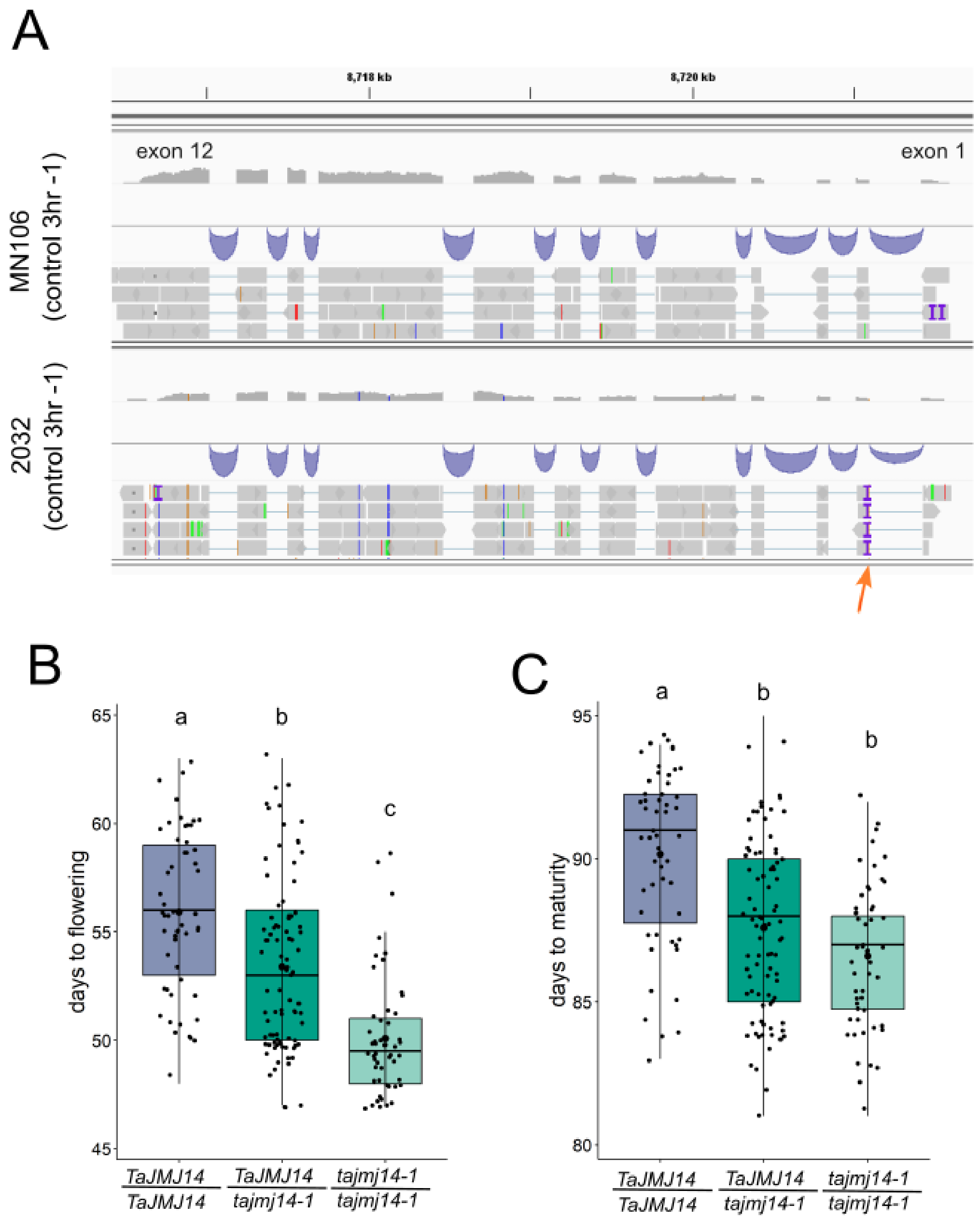


**Supplemental Figure 1. View of *TaJMJ14* transcript read alignment in MN106 and 2032 leaves, and flowering and maturity dates of F2 plants derived from B36 x 2032 cross.** Illumina sequencing reads were mapped to the *T.arvense*_v2 genome, and .bam files of MN106 and 2032 control samples were loaded into IGV (Thorvaldsdóttir et al., 2013). The *TaJMJ14* gene is encoded on the reverse strand of chromosome 7, between positions 8,716,443-8,721,689. The orange arrow highlights earliness marker: an insertion (purple letter I) in the predicted 5'UTR of 2032 samples that was used for genotyping the 2032 (*tajmj14-1*) allele. **(B-C)** F2 plants from a cross of B36 x 2032 were genotyped for the presence of the *tajmj14-1* allele. The days from germination to the appearance of the first flower **(B)** and to the first mature pod **(C)** were recorded. Each data point is an individual F2 plant. Bars = SD; groups indicated by the same letter are not significantly different as assessed by Kruskal-Wallis with Dunn's post-hoc test.

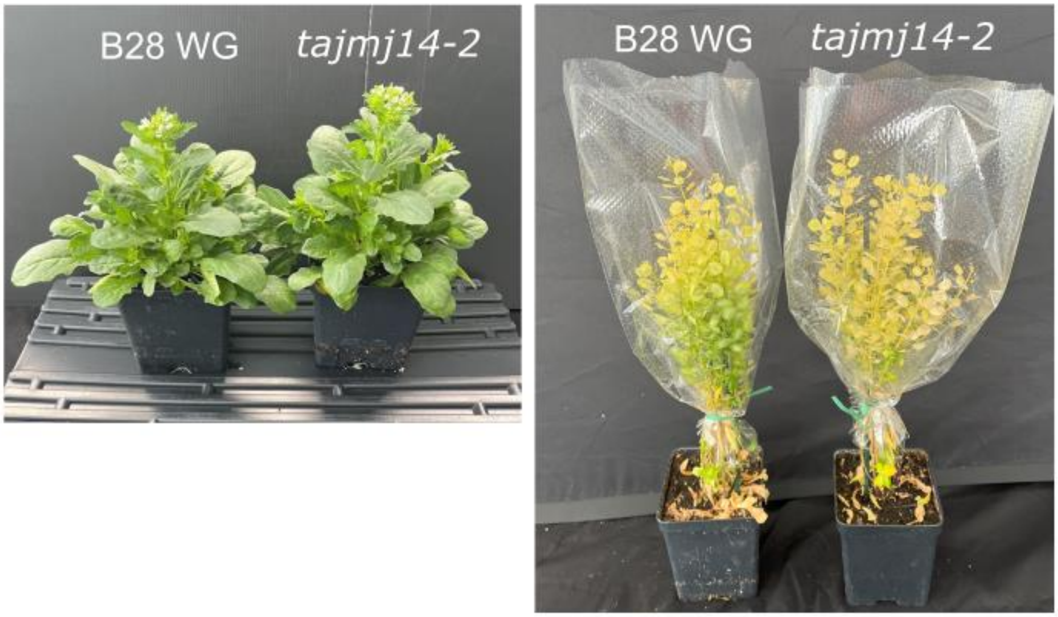


**Supplemental Figure 2. Whole plant phenotypes of *tajmj14-2* T3 and parental B28 WG lines.** Pictures of greenhouse-grown plants during flowering (left, 62 days after germination) and maturity (right, 108 days after germination) windows.

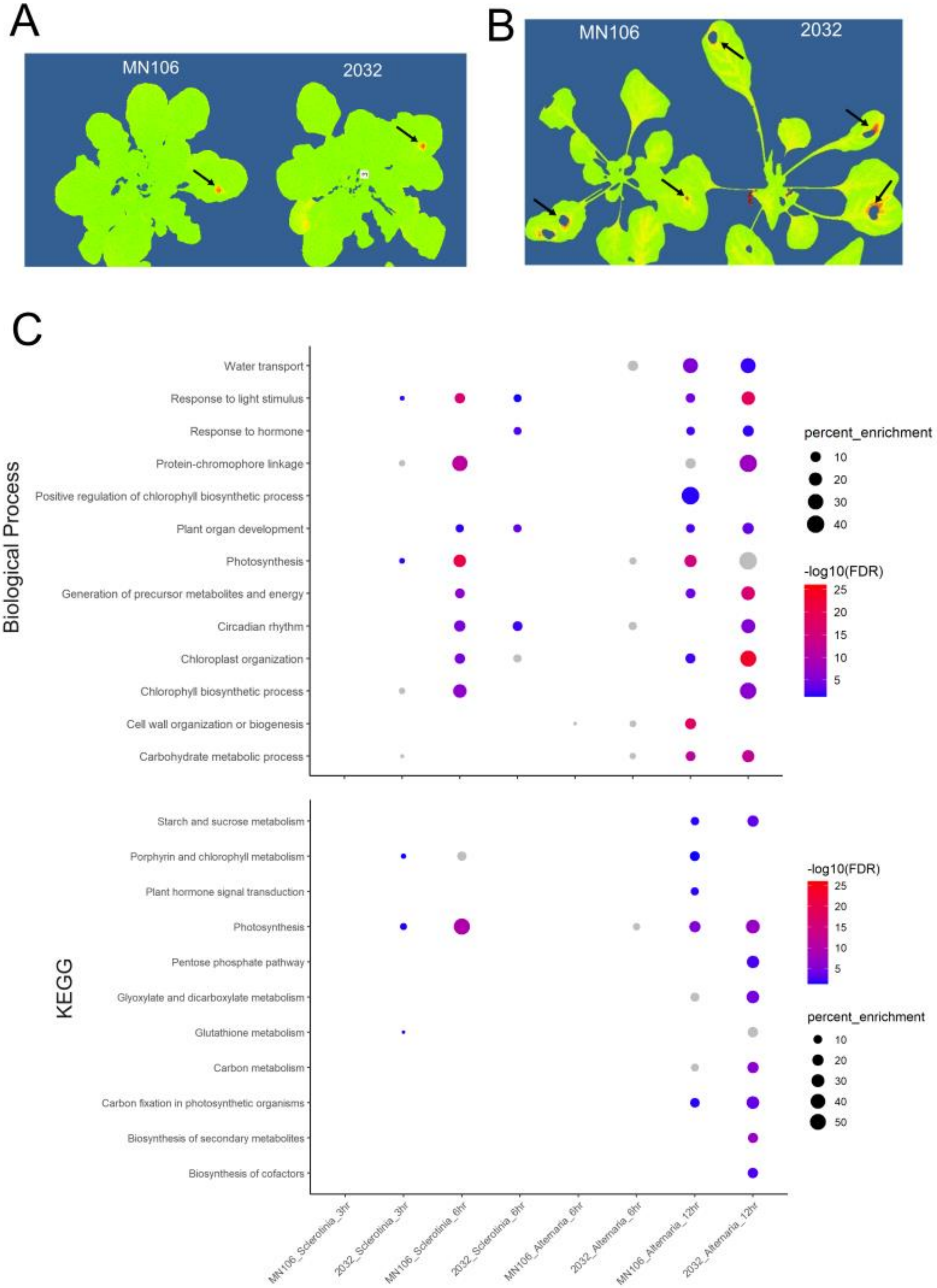
A
MN106
2032
B
MN106
2032
C
Biological Process
Water transport
Response to light stimulus
Response to hormone
Protein-chromophore linkage
Positive regulation of chlorophyll biosynthetic process
Plant organ development
Photosynthesis
Generation of precursor metabolites and energy
Circadian rhythm
Chloroplast organization
Chlorophyll biosynthetic process
Cell wall organization or biogenesis
Carbohydrate metabolic process
percent_enrichment
10
20
30
40
-log10(FDR)
25
20
15
10
5
KEGG
Starch and sucrose metabolism
Porphyrin and chlorophyll metabolism
Plant hormone signal transduction
Photosynthesis
Pentose phosphate pathway
Glyoxylate and dicarboxylate metabolism
Glutathione metabolism
Carbon metabolism
Carbon fixation in photosynthetic organisms
Biosynthesis of secondary metabolites
Biosynthesis of cofactors
-log10(FDR)
25
20
15
10
5
percent_enrichment
10
20
30
40
50
MN106_Sclerotinia_3hr
2032_Sclerotinia_3hr
MN106_Sclerotinia_6hr
2032_Sclerotinia_6hr
MN106_Alternaria_6hr
2032_Alternaria_6hr
MN106_Alternaria_12hr
2032_Alternaria_12hr

**Supplemental Figure 3. Earliest detectable necrosis on *S. sclerotiorum* and *A. japonica*-infected plants and GO enrichment analysis of genes downregulated by infection.** Leaves were inoculated with 6 mm agar plugs of *S. sclerotiorum* for 12 hpi **(A)** or *A. japonica* for 24 hpi **(B)**, and Fv/Fm images were taken using the CropReporter system. The red lesions indicated with black arrows are regions of low photosynthetic efficiency caused by necrotrophic fungal pathogens (Djami-Tchatchou et al., 2023). The holes within the *A. japonica* lesions are an imaging and masking artifact caused by the black color of *A. japonica* agar plugs. **(C)** DEGs downregulated by each infection relative to their own controls were subjected to gene ontology analysis. Presented are the most enriched biological processes (top) and pathways (bottom). Dots are grey if the p-value cutoff (FDR) is > 0.05 and are absent if that pathway is not enriched in that infection.

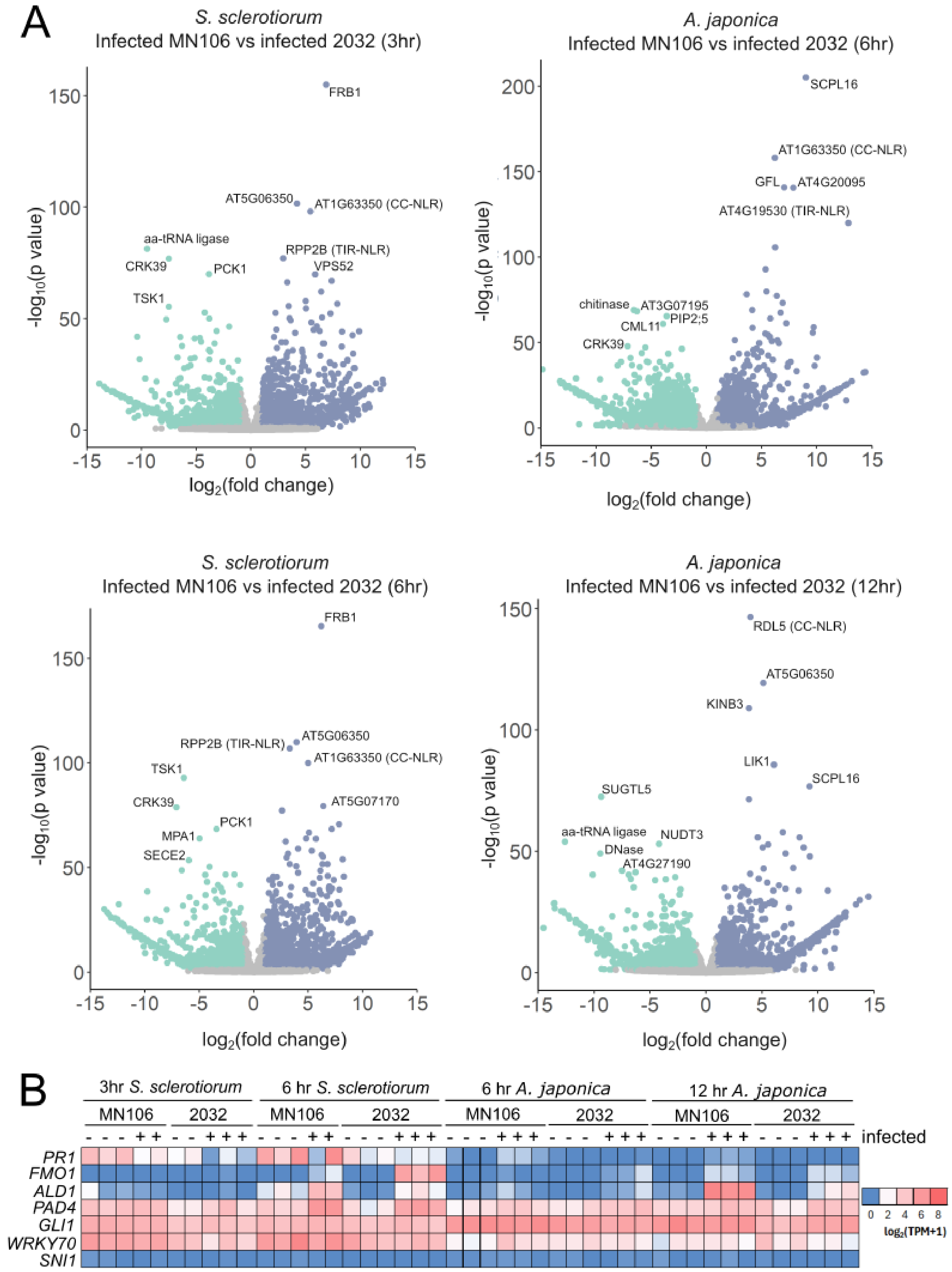
A
S. sclerotiorum
Infected MN106 vs infected 2032 (3hr)
FRB1
AT5G06350
AT1G63350 (CC-NLR)
aa-tRNA ligase
RPP2B (TIR-NLR)
CRK39
PCK1
VPS52
TSK1
A. japonica
Infected MN106 vs infected 2032 (6hr)
SCPL16
AT1G63350 (CC-NLR)
GFL
AT4G20095
AT4G19530 (TIR-NLR)
chitinase
AT3G07195
CML11
PIP2;5
CRK39
S. sclerotiorum
Infected MN106 vs infected 2032 (6hr)
FRB1
RPP2B (TIR-NLR)
AT5G06350
AT1G63350 (CC-NLR)
TSK1
CRK39
AT5G07170
PCK1
MPA1
SECE2
A. japonica
Infected MN106 vs infected 2032 (12hr)
RDL5 (CC-NLR)
AT5G06350
KINB3
LIK1
SCPL16
SUGTL5
aa-tRNA ligase
NUDT3
DNase
AT4G27190
$-log_{10}$(p value)
$log_2$(fold change)
B
3hr S. sclerotiorum
6 hr S. sclerotiorum
6 hr A. japonica
12 hr A. japonica
MN106
2032
infected
PR1
FMO1
ALD1
PAD4
GLI1
WRKY70
SNI1
0 2 4 6 8
$log_2$(TPM+1)

**Supplemental Figure 4. Most-enriched genes upon infection in MN106 and 2032 plants, and expression patterns of AtJMJ14-regulated genes in infected pennycress. (A)** Volcano plots showing the abundance of transcripts enriched in infected MN106 (right) compared to infected 2032 (left) samples for each infection. Gray dots are those transcripts that did not meet thresholds of $\log_2$(fold change) > 2 and p-value < 0.05. The 5 most enriched transcripts for each genotype are labelled. **(B)** Expression of selected genes whose expression was regulated by AtJMJ14 in Li et al. (2020). Expression is displayed in $\log_2$(transcripts per million +1) form.

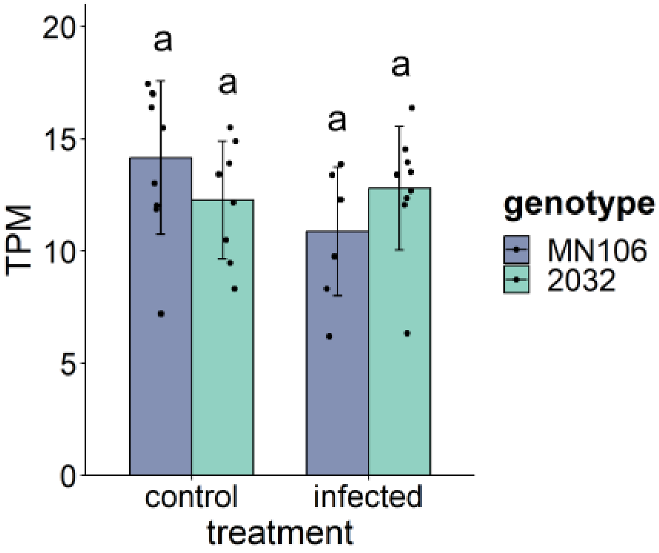


**Supplemental Figure 5. Expression of *TaJMJ14* transcript in control and infected leaves.** RNAseq reads aligning to the genomic coordinates of *TaJMJ14* (Chr7:8,716,443-8,721,689) expressed in transcripts per million (TPM). Groups indicated with the same letter are not significantly different according to One-Way ANOVA with Scheffe's post-hoc test.

**Supplemental Table 1. List of primers utilized in this study.**

| Primer name | Purpose | sequence (5' - 3') |
|---|---|---|
| JMJ14_PS1_For | Sanger genotyping of editing by JMJ14 PS1 | CGGTTGATGACGCTCCCATA |
| JMJ14_PS1_Rev | Sanger genotyping of editing by JMJ14 PS1 | TTGCTTGCTCAACTATCCGC |
| JMJ14_PS2_For | Sanger genotyping of editing by JMJ14 PS2 | GAATATTGGCGGATAGTTGAGCA |
| JMJ14_PS2_Rev | Sanger genotyping of editing by JMJ14 PS2 | CATTGCGTTTTCAAAGGACTCAG |
| JMJ14_PACE_WT_HEX | PACE genotyping of *jmj14-1* allele | GAAGGTGACCAAGTTCATGCTGTCGAGGGTTAAAAGTACCGGGT |
| JMJ14_PACE_mutant_FAM | PACE genotyping of *jmj14-1* allele | GAAGGTCGGAGTCAACGGATTCGAGGGTTAAAAGTACCGGGC |
| JMJ14_PACE_common | PACE genotyping of *jmj14-1* allele | CCCTTCCATTTACAGTGAATTTAGGGTTT |
| Ajs_ITS_For | qPCR detection of *A. japonica ITS* DNA | GGCGTCTTGTCTCCAGTTTG |
| Ajs_ITS_Rev | qPCR detection of *A. japonica ITS* DNA | CCTACCTGATCCGAGGTCAA |
| TaAct_qPCR_For | qPCR detection of *T. arvense actin* DNA | GGAATCCACGAGACGACCTA |
| TaAct_qPCR_Rev | qPCR detection of *T. arvense actin* DNA | CTTGGTGCAAGTGCTGTGAT |

**Supplemental File 1.** Genomic and coding sequence (CDS) sequences of *TaJMJ14* from MN106 and 2032 in FASTA format. Clustal Omega alignment of genomic *TaJMJ14* sequences. Clustal Omega alignment of *TaJMJ14* genomic sequences from pennycress pangenome (https://phytozome-next.jgi.doe.gov/pennypan/).

**Supplemental File 2.** Differentially expressed genes (DEGs) from RNAseq present in all four infection conditions (2 pathogens, 2 time points each) tested. Tab 1: DEGs more highly expressed in infected 2032 leaves compared to infected MN106 leaves. Tab 2: DEGs more highly expressed in infected MN106 leaves compared to 2032 leaves (tab 2).
